# Biodegradable silica nanoparticles for efficient linear DNA gene delivery

**DOI:** 10.1101/2023.12.05.569925

**Authors:** Andrés Ramos-Valle, Henning Kirst, Mónica L. Fanarraga

## Abstract

Targeting, safety, scalability, and storage stability of vectors are still challenges in the field of nucleic acid delivery for gene therapy. Silica-based nanoparticles have been widely studied as gene carriers, exhibiting key features such as biocompatibility, simplistic synthesis and enabling easy surface modifications for targeting. However, the ability of the formulation to incorporate DNA is limited, which restricts the number of DNA molecules that can be incorporated into the particle, thereby reducing gene expression.

Here we use polymerase chain reaction (PCR)-generated linear DNA molecules to augment the coding sequences of gene-carrying nanoparticles, thereby maximizing nucleic acid loading and minimizing the size of these nanocarriers. This approach results in a remarkable 16-fold increase in protein expression six days post-transfection in cells transfected with particles carrying the linear DNA compared with particles bearing circular plasmid DNA. The study also showed that the use of linear DNA entrapped in DNA@SiO_2_ resulted in a much more efficient level of gene expression compared to standard transfection reagents. The system developed in this study features simplicity, scalability, and increased transfection efficiency and gene expression over existing approaches, enabled by improved embedment capabilities for linear DNA, compared to conventional methods such as lipids or polymers, which generally show greater transfection efficiency with plasmid DNA. Therefore, this novel methodology can find applications not only in gene therapy but also in research settings for high throughput gene expression screenings.

## 1 Introduction

Gene therapy has made significant progress in recent years, leading to the approval of several groundbreaking therapies that have paved the way for the treatment of several diseases, with remarkable benefits for patients.[1,2] This technology includes genome editing,[3] production of therapeutic proteins in damaged areas,[4] or boosting the body’s natural defense mechanisms against diseases.[5] These approaches can be used as therapies to mitigate the impact of acquired mutations or enhance cellular functions affected by diseases. Additionally, gene transfer allows disease modeling in mice and other species, facilitating the exploration of therapeutic approaches and addressing scientific questions that are challenging to investigate in humans.[6] The technology has also significant potential in advancing agriculture by increasing food security, reducing the need for pesticides, and improving environmental sustainability through genetic modification.[7,8]

An effective strategy for gene transfer is to use viral vectors such as retroviruses, lentiviruses, adenoviruses, and adeno-associated viruses.[9] Among them, lentiviral vectors are commonly used because they enable *ex vivo* cell modification before patient reintroduction.[10] However, the use of viral vectors carries certain risks. One concern is the potential for introducing random DNA into the host genome, which can compromise the safety and efficacy of the treatment.[11,12] Hence, several critical issues need to be addressed, including vector toxicity, non-specificity, and instability for the clinical validation of *in vivo* gene transfer methods for clinical use.

Significant advances in vector technology, particularly synthetic nucleic acid vectors, offer promising gene delivery strategies. Among these, nucleic acid-loaded nanoparticles serve as valuable tools for efficient cargo delivery, enabling gene expression,[13] transient protein expression,[14] and genome editing in target cells.[15] Liposomes are among the most promising nano-vectors due to their customizability, ease of preparation, ability to accommodate genes of varying sizes, and have a low immunogenicity or toxicity.[16,17] Solid-lipid nanoparticle formulations loaded with nucleic acids have also shown potential as delivery vehicles for nucleic acids in cancer treatment. However, even though lipid-based formulations have the potential to overcome different barriers, their stability remains a concern.[18] Furthermore, transfection efficiency with linear DNA using lipid based nanocarriers is disappointingly low. Also, compared to viruses, lipid-based nano-delivery systems are challenging to target *in vivo* because they tend to concentrate in the liver or spleen.[19] Besides, they often experience a rapid release of encapsulated DNA, which results in a short burst of recombinant gene expression, which may compromise the viability of transfected cells.[20] Despite varying successrates in *in vitro* and preclinical research, with some gene delivery systems undergoing validation as prospective therapeutics, the development of efficient and effective gene delivery vectors continues to face challenges. Safety concerns with viral or polymer-based delivery systems remain an issue, as do the high cost and scalability barriers of complex nanoparticle synthesis.[21]

Silica particles (SiO_2_) have emerged as highly valuable systems for delivering drugs and nucleic acids.[22] These nanoscale vehicles possess distinct morphologies and structures, providing numerous advantages such as large specific surface areas, adjustable particle and pore sizes, favorable mechanical and thermal stability, low toxicity, and remarkable biocompatibility.[23] Also, the properties and functionality of silica nanoparticles can be tailored by harnessing their surface chemistry,[24] which opens new avenues for their use in nanotechnology. Colloidal silica nanoparticles, in particular, have found diverse applications as additives in the production of food, cosmetics, pharmaceuticals, and other applications.[25–29] As a result, the field of drug delivery has extensively explored silica nanoparticles for over six decades.

Tailoring the size, shape, and surface functionalization of silica materials has led to increased biocompatibility and efficiency of delivery. There are different methods of delivering DNA using silica nanoparticulate systems. One of them involves adsorbing DNA on the nanoparticle surface.[30–32] However, this approach may not provide sufficient protection for the DNA and limits DNA loading capacity, negatively impacting transfection and gene expression efficiencies. To address this issue, a newer design has been developed, where the DNA is directly embedded within the formulation of the nanoparticle itself.[33] This approach ensures better protection of the nucleic acids and enhances their delivery efficiency, providing a protective barrier against environmental nucleases or reactive oxygen species (ROS) and enhancing the stability of the genetic cargo. Once captured by a cell, the DNA@SiO_2_ particles dissolve intracellularly, release the encapsulated DNA triggering adequate cell transfection rates.[33] While silica-based nanosystems have great potential as gene delivery systems due to their targeting, safety, scalability, and storage stability, the actual volume of the silica particles limits DNA capacity and transgene expression. And although larger particles could be synthesized to encapsulate more DNA molecules and enhance gene expression, this negatively impacts particle uptake and transfection efficiency. Here, we enhance DNA@SiO_2_ particle gene expression by optimising the process used to synthesise the particles. We have focused on reducing the size of the DNA molecules to essential sequences, thereby increasing the total number of DNA molecules. The expected outcome is the development of a simple, low-cost system that offers a promising alternative to other gene delivery systems that are less versatile and stable.

## 2 Material and methods

### 2.1 DNA production and purification

The p-DNA was produced using standard procedures in *E. coli* DH5α that had been transformed with the different plasmids used in the study (Figure S1). Bacteria were cultured overnight in Luria-Bertani (LB) broth medium. DNA was extracted and purified using a kit (PureLink™ HiPurePlasmid Maxiprep, Thermo Fisher) following the manufacturer’s protocol. DNA was precipitated and resuspended in ddH_2_O at 2.5 µg/ml for direct use in the particle synthesis.

Linear DNA was produced using the PCR touchdown technique with the corresponding p-DNA as a template (Figure S1). The PCR mixture was composed of forward and backward-designed primers (2.5 µM) purchased from IDT, Master Mix NZYProof 2 × (1/2 of total PCR volume) containing Taq II Polymerase Enzyme, dNTPs, buffers, and plasmid DNA template (100 ng / 50 µl of total volume), (Table S1).

### 2.2 Synthesis and characterization of DNA@SiO_2_

Silica spheres containing DNA were prepared using a modified Stöber method previously described.[34–37] In brief, a mixture of EtOH and 4.5 µg of the corresponding p-DNA (1.4 x 10_-5_ M) previously dispersed in 18 µl of nuclease-free ddH_2_O (10 M) was stirred at 1,200 r.p.m.

Subsequently, ammonia 25 % (1.22 M) and TEOS (0.24 M) were added to this mixture (Table 1). The reaction mixture was then vortexed for 2 h at 1,200 r.p.m. Silica spheres were centrifuged, washed, and stored in EtOH. The concentration of the final components in the mixture considers EtOH as a solvent and is expressed in molarity (M).

**Table 1.** Optimization of DNA@SiO_2_ particle synthesis.

| Entry | Synthesis |  |  |  |  | Characterization |  |  |
| --- | --- | --- | --- | --- | --- | --- | --- | --- |
|  | NH<sub>3</sub><br>/M | TEO<br>S/ M | H<sub>2</sub>O<br>/M | DNA |  | Size<br>(TEM) | Z-Size<br>(DLS) | PDI<br>(DLS) |
|  |  |  |  | Shape | [DNA] / M<br>(µg DNA/mg SiO<sub>2</sub>) |  |  |  |
| type a | 1.06 | 0.24 | 3.32 | plasmid | 1.4 x 10<sup>-5</sup> (4.5 µg) | 678 ± 51 | - | - |
| type b | 1.22 | 0.24 | 9.99 | plasmid | 1.4 x 10<sup>-5</sup> (4.5 µg) | 388 ± 36 | - | - |
| type c | 0.34 | 0.24 | 14.0 | plasmid | 1.4 x 10<sup>-5</sup> (4.5 µg) | 186 ± 15 | - | - |
| p-DNA#1 | 1.22 | 0.24 | 9.99 | plasmid | 1.6 x 10<sup>-6</sup> (0.5 µg) | 468 ± 28 | 601 ± 28 | 0.18 ± 0.08 |
| p-DNA#2 | 1.22 | 0.24 | 9.99 | plasmid | 4.8 x 10<sup>-6</sup> (1.5 µg) | 274 ± 19 | 358 ± 18 | 0.2 ± 0.1 |
| p-DNA#3 | 1.22 | 0.24 | 9.99 | plasmid | 9.6 x 10<sup>-6</sup> (3.0 µg) | 262 ± 16 | 324 ± 3 | 0.26 ± 0.02 |
| p-DNA#4 | 1.22 | 0.24 | 9.99 | plasmid | 1.4 x 10<sup>-5</sup> (4.5 µg) | 393 ± 22 | 413 ± 67 | 0.23 ± 0.06 |
| p-DNA#5 | 1.22 | 0.24 | 9.99 | plasmid | 2.9 x 10<sup>-5</sup> (9 µg) | * | 1874 ± 25 | 0.99 ± 0.05 |
| p-DNA#6 | 1.22 | 0.24 | 9.99 | plasmid | 4.8 x 10<sup>-5</sup> (15 µg) | * | 3770 ± 17 | 1.5 ± 0.2 |
| I-DNA#1 | 1.22 | 0.24 | 9.99 | linear | 1.6 x 10<sup>-6</sup> (0.5 µg) | 384 ± 40 | 325 ± 12 | 0.41 ± 0.02 |
| I-DNA#2 | 1.22 | 0.24 | 9.99 | linear | 4.8 x 10<sup>-6</sup> (1.5 µg) | 311 ± 18 | 314 ± 28 | 0.35 ± 0.03 |
| I-DNA#3 | 1.22 | 0.24 | 9.99 | linear | 9.6 x 10<sup>-6</sup> (3.0 µg) | 284 ± 20 | 236 ± 60 | 0.36 ± 0.06 |
| I-DNA#4 | 1.22 | 0.24 | 9.99 | linear | 1.4 x 10<sup>-5</sup> (4.5 µg) | 260 ± 22 | 250 ± 28 | 0.33 ± 0.03 |
| I-DNA#5 | 1.22 | 0.24 | 9.99 | linear | 2.9 x 10<sup>-5</sup> (9 µg) | * | 1034 ± 30 | 0.7 ± 0.3 |

**Notes:** Effect of different concentrations of the Stöber mixture (a-c). Effects of different amounts of DNA (plasmid and linear) on particle size and dispersity. Figures S1-S11 show the TEM and DLS results of characterization. Asterisk in places where (\*) TEM sizes could not be estimated due to aggregates (see Figures S5-S6, Figures S11).

Silica spheres containing linear DNA were prepared using a similar method. In a typical experiment, a mixture of, EtOH and 4.5 µg of the corresponding l-DNA (1.4 x 10_-5_ M) previously dispersed in 18 µl of nuclease-free ddH_2_O (10 M) was stirred at 1,200 r.p.m. for 5 min. Subsequently, ammonia 25 % (1.22 M) and TEOS (0.24 M) were added to this mixture (Table 1). The reaction mixture was then vortexed for 2 h. Finally, silica spheres were separated from the suspension by centrifugation at 6,500 r.p.m. for 5 min and washed twice with EtOH. The particles were then washed repeatedly. Particles were dispersed in absolute EtOH (0.5 mg/ml) and 10 µl was applied to the corresponding carbon film on a copper grid for Transmission Electron Microscopy (TEM) analysis. The samples were dried for 1 h at 25 °C and subsequently introduced into a microscope sampler. TEM images were obtained with a JEM1011 TEM equipped with a high-resolution Gatan digital camera (JEOL, Japan). The images were then analyzed using ImageJ software. DLS size distribution measurements of p-DNA#1-6 and l-DNA#1-5 were carried out on a Malvern Ultra Zetasizer at 25 ºC using filtered dH_2_O as a dispersant. The measurements were repeated three times for each sample, and the data are presented as mean ± standard deviation (SD).

### 2.3 Cell lines

HEK 293T (Human Embryonic Kidney) cells were maintained in Iscove’s modified Dulbecco’s medium (IMDM) supplemented with 10% fetal bovine serum and antibiotics in a 5% CO2 incubator at 37o C.

### 2.4 Cell transfection assays

HEK 293T cells were reseeded in a 24-well plate before transfection. A 100 µg NPs/ml DNA@SiO_2_ particle resuspended in culture medium was added to each well. The medium was replaced after 16 hours. For LipofectamineTM 2000 transfection of HEK 293T cells, 3.13 µl of reagent was diluted in 496.85 µl of serum-free IMDM medium and incubated for 5 minutes. 2.5 µg of p-DNA was diluted in 500 µl of serum-free IMDM according to the manufacturer’s instructions. Cells were fixed in a 4% PFA solution for 15 min before qualitative or quantitative characterization. Cells were stained with DAPI, washed in PBS, mounted on crystal glass coverslips, and imaged using a Nikon AIR confocal microscope. Flow cytometry was performed on fixed transfected cells. Approximately 10,000 cells were analyzed using a CytoFLEX (Beckman Coulter) equipped with 3 excitation lasers (488 nm x 50mW; 638 nm x 50mW; 405 nm x 80mW) and 13 fluorescence detectors. Data corresponding to the positive cell distribution were obtained with the appropriate single-cell gating. Three different replicates were performed for each experiment. Data are expressed as mean ± standard deviation (SD).

### 2.5 Cell viability assays

HEK 293T cells were reseeded into 96-well culture plates at a density of 2 x 10_4_ cells/well and incubated for 24 h. DNA@SiO_2_ particles were washed in Phosphate Buffer Saline (PBS) and were diluted with Iscove’s Modified Dulbecco′s Medium (IMDM) with sera. Mixtures were added at concentrations ranging from 0 to 400 µg/ml (see Figure 3). After incubation at 37 _o_C for 16 h, the medium in each well was renewed, and the cells were incubated for 24 h and 72 h before the addition of 10 μl of the MTT labeling reagent (final concentration 0.5 mg/ml). After incubation for 4 h in a humidified atmosphere (37 °C, 5-6.5% CO_2_), 100 μl of the solubilization solution was added to each well. The plates were allowed to stand overnight in the incubator in a humidified atmosphere. The absorbance of the samples was measured using a microplate reader.

### 2.6 Statistical analysis

A total of 10,000 cells were analyzed per condition in three replicas of each experiment in Figures 1, 2, 3, and 4. Analysis *t-*tests were employed to compare the Mean Fluorescence Intensities (related bars), and 0.05 statistical analysis was used. Error bars represent SD.

## 3. Results

### 3.1 Excess plasmid DNA results in aberrant DNA@SiO_2_ particles

Aiming to reduce the DNA@SiO_2_ particle size, we conducted a series of experiments, varying the stoichiometric ratios of NH_3_, H_2_O, and TEOS in the Stöber method, while maintaining consistent amounts of plasmid DNA (p-DNA) per particle. This allowed us to synthesize DNA@SiO_2_ particles with sizes ranging from 100 to 620 nm, labeled as type a-c (Table 1, Figures S2-S12). Due to the considerable size variation observed in type-c particles and their negative impact on cellular uptake, we narrowed our focus to optimize type-b particles, with a diameter of approximately 350 nm.

To enhance transfection efficiency and achieve higher levels of protein expression, we proceeded to assess how the DNA content in the formulation impacts nanoparticle characteristics, such as size, morphology, and polydispersity. Our goal was to identify strategies that can optimize particle properties and maximize nucleic acid loading, thereby improving overall transfection efficiency. Proof-of-concept experiments were performed utilizing plasmid DNA (p-DNA) encoding the the histone H2B tagged with yellow fluorescent protein (H2B:YFP) leading to fluorescent nuclei after transfection and protein expression. Various concentrations ranging between 1.6 x 10_-6_ M and 4.8 x 10_-5_ M of DNA were chosen to investigate the impact of nucleic acid concentration on the properties of the particles. Interestingly, we observed that with an increasing amount of p-DNA per particle the particle size decreased slightly (Table 1). The sizes ranged from 600 nm for p-DNA#1 to 350 nm for p-DNA#3 (Figure 1). However, at higher DNA concentrations, there was a significant increase in particle polydispersity that was likely due to the formation of silica aggregates in p-DNA#5-6 (Figures 1c, S5, S6). At a concentration of 4.5 µg p-DNA per mg SiO_2,_ the particles demonstrated the best balance of polydispersity, size, and DNA loading. This concentration yielded optimal results in terms of achieving a desirable particle size distribution and loading capacity while minimizing polydispersity.

**Figure 1.**
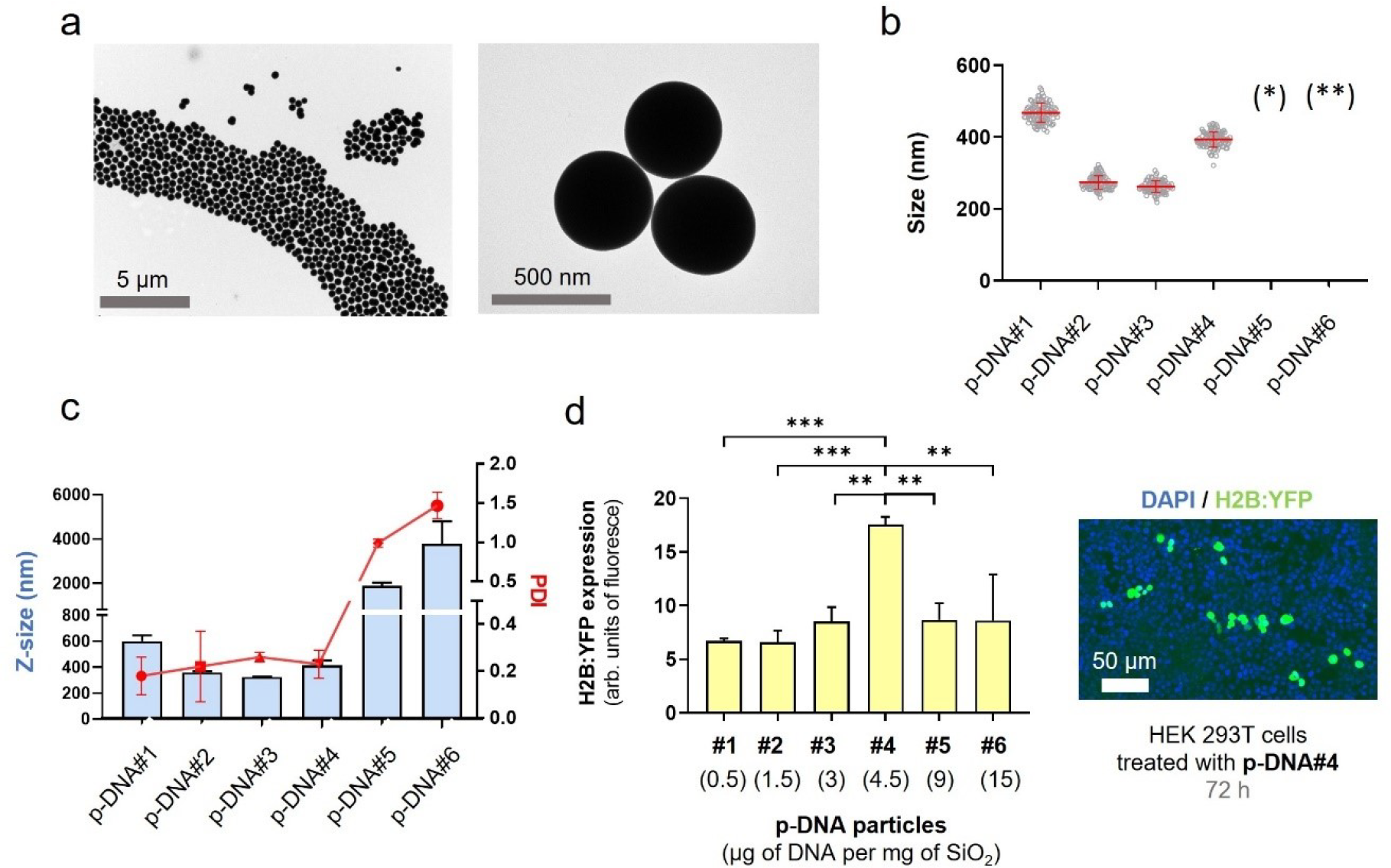
p-DNA@SiO_2_ particles. **(a)** TEM images of p-DNA#4 particles with 4.5 µg of plasmid H2B:YFP DNA. **(b)** Particle diameter (mean ± SD) measured by TEM within a scatter dot plot comparing p-DNA#1-6 (*n* = 100). Values for p-DNA#5 and p-DNA#6 are not included due to the presence of aggregates (see Figures S8-S9). (*)(**) p-DNA#5-6 not included due to the presence of aggregates (see Figures S5-S6). **(c)** DLS characterization of p-DNA#1-6 bar chart of Z-size (mean ± SD, left Y axis, *n* = 3) and dot plot in red showing the polydispersity index (PDI) (mean ± SD, right Y axis, *n* = 3). **(d)** Flow cytometry quantification of the transfection efficiency (mean fluorescence intensity) of the p-DNA#4 DNA@SiO_2_ particles and a representative confocal microscopy image of HEK 293T cells expression the recombinant H2B:YFP protein 72 h after transfection. Data are shown as the mean ± SD of 3 experimental replicas (*n =* 10,000 cells/replica, *t-*test, \**p* < 0.05, and \*\*\**p* < 0.001).

Next, we evaluated the efficiency of various particle types on transfection using HEK 293T cells. This revealed that protein expression levels for particle types p-DNA#1-4 increased gradually and proportionally with higher amounts of p-DNA (Figure 1d). Our results show (Table 1, Figure 1), that a balance is required between the amount of DNA and the concentration of the Stöber components to permit silanol polymerization and effective DNA loading while ensuring minimal polydispersity and efficient transfection of synthesized DNA@SiO_2_ particles. The particles p-DNA#4 are the most suitable candidates because they show a well-balanced combination of maximal gene cargo loading, optimal particle morphology, and transfection efficiency (Figure 1). Furthermore, our experiments show that the total amount of DNA integrated into a silica matrix acts as a limiting factor to ensure the preservation of acceptable particle size and polydispersity index (PDI) values.

### 3.2 Linearization of plasmid DNA: influence of l-DNA size, shape and mass on DNA@SiO_2_ particle synthesis

With this understanding, we next aimed to enhance gene delivery by increasing the coding DNA (gene copy numbers) while maintaining the total DNA mass in the formulation. To achieve this, we used PCR with specific primers to selectively eliminate elements non-essential for transfection of human cells, such as the origin of replication, prokaryotic selection marker, and multiple cloning sites. Through this approach, we successfully generated linear DNA (l-DNA) that exclusively contained the essential DNA sequence necessary for efficient gene expression in human cells. The synthesized l-DNA consisted of three essential elements: (i) a promoter sequence including the ribosome binding site for gene expression, (ii) the coding sequence of the gene, and (iii) a transcriptional terminator. By using l-DNA instead of p-DNA, we were able to reduce the size of the DNA molecule by approximately 60% (Figure S12).

After obtaining the PCR products, we quantified them by UV-VIS 280 nm absorbance and confirmed their correct size through electrophoresis. To assess their functionality and ability to induce gene expression before silica encapsulation, we tested the l-DNA molecules generated by PCR through transfection into mammalian cells using Lipofectamine™ 2000. Transfections performed using conventional methods based on lipid vectors exhibited a notable reduction in efficiency compared to transfections using p-DNA consistend with previous observations.[38] Having confirmed the functionality of the PCR-generated l-DNA molecules, we incorporated them into the synthesis reaction at different concentrations, ranging from 0.5 to 9 μg per mg SiO_2_ (Table 1). Similar to the observations with p-DNA, l-DNA quantities exceeding 4.5 μg led to particle aggregation, resulting in Z-sizes of approximately 1000 nm and a PDI of 0.7 ± 0.3 (Figures 2c, S11). Nonetheless, particles loaded with 4.5 μg of l-DNA (l-DNA#4) demonstrated an ideal size range of 240-280 nm in diameter with a monodisperse distribution. These results indicate that, regardless of the shape and size of the DNA (p-DNA or l-DNA), the total mass of DNA in the Stöber mixture is limited to 4.5 μg per mg SiO_2_ to yield acceptable nanoparticle properties. Functional tests on cells showed that l-DNA@SiO_2_ particles exhibited enhanced gene transfection efficiency compared to p-DNA@SiO_2_ particles (Figure 2d), indicating that our strategy of increasing the amount of DNA molecules by encapsulating l-DNA is suitable.

**Figure 2.**
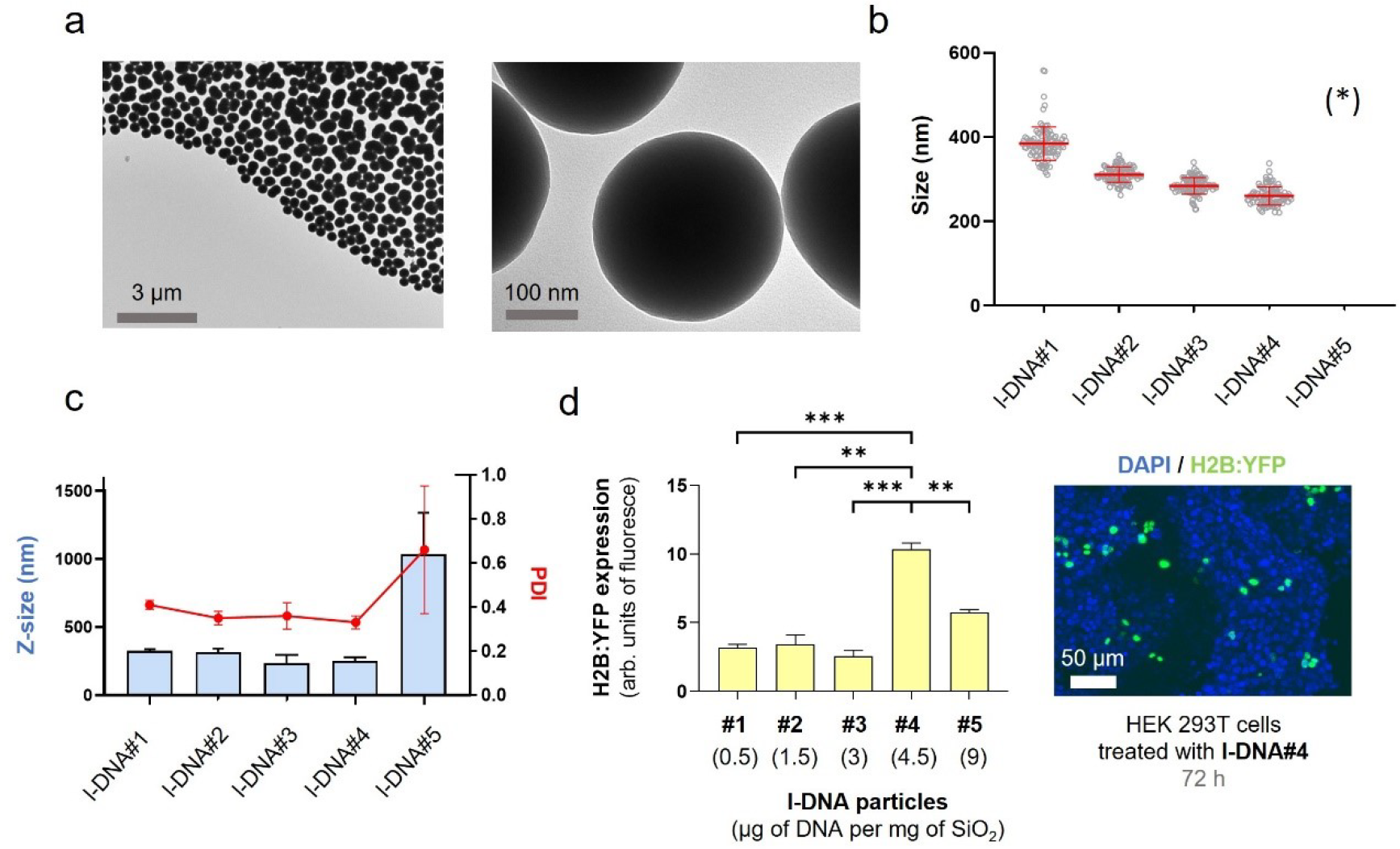
l-DNA@SiO_2_ particles. **(a)** TEM images of particles l-DNA#4 with 4.5 µg of linear H2B:YFP DNA. (**b)** Particle diameter (mean ± SD) measured by TEM within a scatter dot plot comparing l-DNA#1-5 (*n* = 100). (*) l-DNA#5 is not included due to the presence of aggregates (see Figure S11). **(c)** l-DNA#1-6 bar chart of Z-size (mean ± SD, left Y axis, *n* = 3) and dot plot in red showing Polydispersity index (PDI) (mean ± SD, right Y axis, *n* = 3). **(d)** Flow cytometry quantification of the transfection efficiency (mean fluorescence intensity) of the l-DNA#4 DNA@SiO_2_ particles and a representative confocal microscopy image of HEK 293T cells expression the recombinant H2B:YFP protein 72 h after transfection. Data are shown as the mean ± SD of 3 experimental replicas (*n =* 10,000 cells/replica, *t-*test, \**p* < 0.05, and \*\*\**p* < 0.001).

### 3.3 DNA@SiO_2_ particles improve transfection efficiency of linear DNA over plasmid DNA

The transfection efficiency of l-DNA@SiO_2_ particles with different amounts of l-DNA was evaluated for comparison with p-DNA particles. Specifically, we prepared particles with a consistent number of DNA molecules, equivalent to 1.5 µg DNA per mg SiO_2_ (referred to as l-DNA#2). Furthermore, taking advantage of the capability of l-DNA to load approximately three times more DNA molecules compared to p-DNA, we encapsulated 4.5 µg of l-DNA per mg of SiO_2_ (referred to as l-DNA#4), ensuring that the total mass of DNA was comparable to that contained in the original p-DNA@SiO_2_ particles (referred to as p-DNA#4).

Quantification and comparison of protein expression 72 hours after transfection using flow cytometry revealed significantly higher levels in both l-DNA#2 and l-DNA#4 particles compared to p-DNA#4 particles, with 8-fold and 16-fold increases, respectively (Figure 3a). On day 6 post-transfection, the expression of l-DNA was still higher than p-DNA, beeingy twice as high as that of p-DNA, while on days 10 and 14 there were no significant differences between the two. We propose that this pattern of protein expression is due to the process of silica dissolution, which is accelerated by the intracellular presence of monovalent cations. [33,39] The entrapment of each l-DNA molecule inside SiO_2_ spheres would be composed of less amount of silica due to their decreased size compared to p-DNA. Assuming a constant SiO_2_ dissolution rate in physiological conditions, l-DNA molecules would be faster released than larger plasmid templates (since they required a complete dissolution of their individual silica entrapment). This hypothesis would explain a faster expression with l-DNA#2 particles than p-DNA#4 (Figure 3.b).

**Figure 3.**
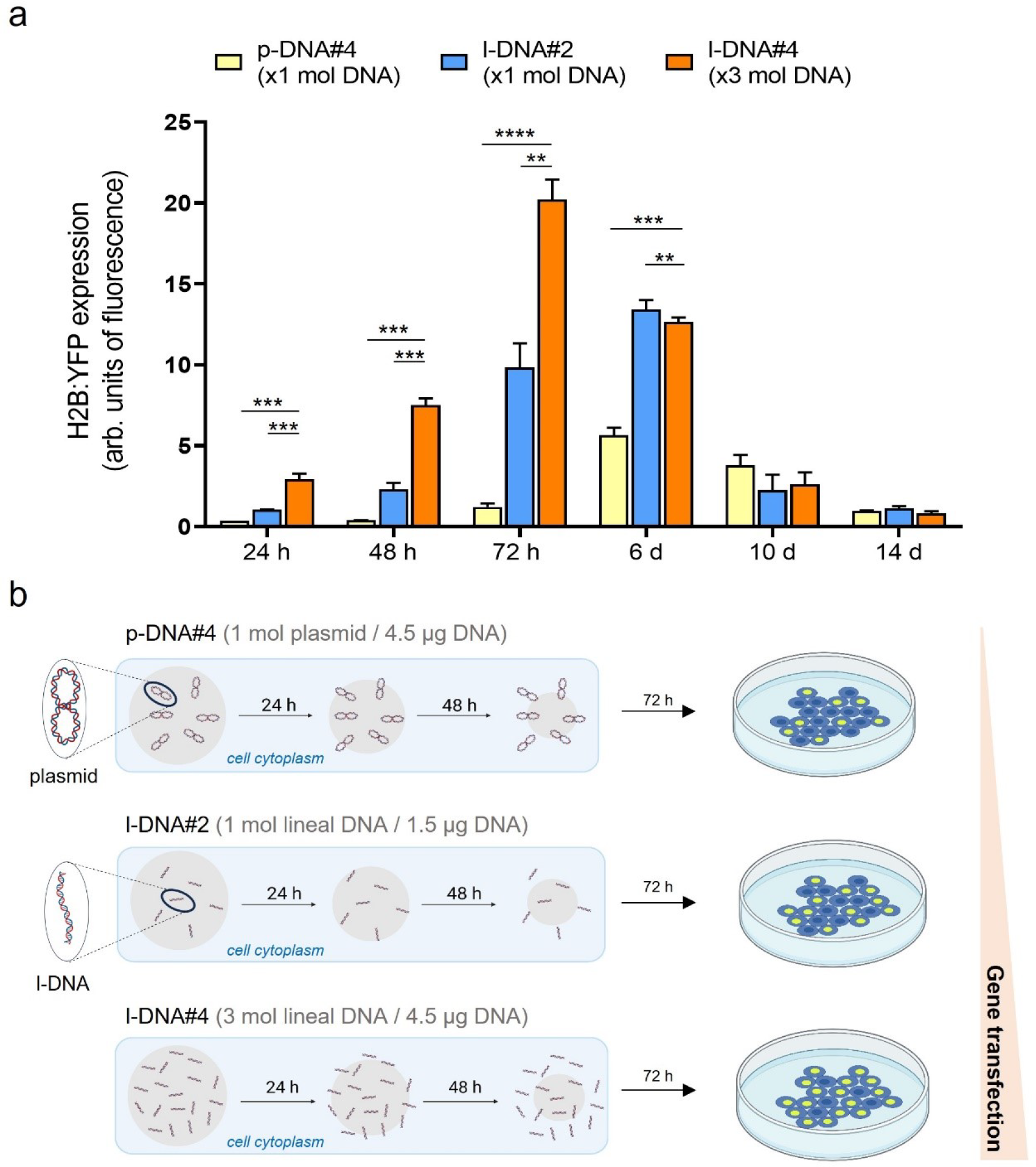
**(a)** Quantification of the transfection efficiency (mean fluorescence intensity) using p-DNA#4, l-DNA#2, and l-DNA#4 particles. Data are shown as the mean ± SD of 3 experimental replicas (*n* = 10,000 cells/replica, *t-*test, ** *p* < 0.01, \*\*\**p* < 0.001, and \*\*\*\**p* < 0.0001). Protein expression profile from 24 h to 14 days. **(b)** Mechanistic proposal of differences in gene transfection efficiency for p-DNA#4, l-DNA#2, and l-DNA#4.

### 3.4 Effect of particle concentration on cell transfection and viability

To find the optimal concentration of l-DNA#4 particles in transfection experiments maximizing gene expression while minimizing cell damage, we incrementally increased the nanoparticle concentration in the culture. We assessed the changes in recombinant gene expression levels and also examined the cytotoxicity resulting from the presence of these nanoparticles in the medium at both 24 and 72 hours. Gene expression quantification was performed using flow cytometry, while toxicity was evaluated through an MTT analysis. Our findings show that using l-DNA#4 particles at concentrations ranging from 100 to 200 µg DNA/ml resulted in the most effective delivery of l-DNA, showcasing promising applications in the field of gene therapy (Figure 4).

**Figure 4.**
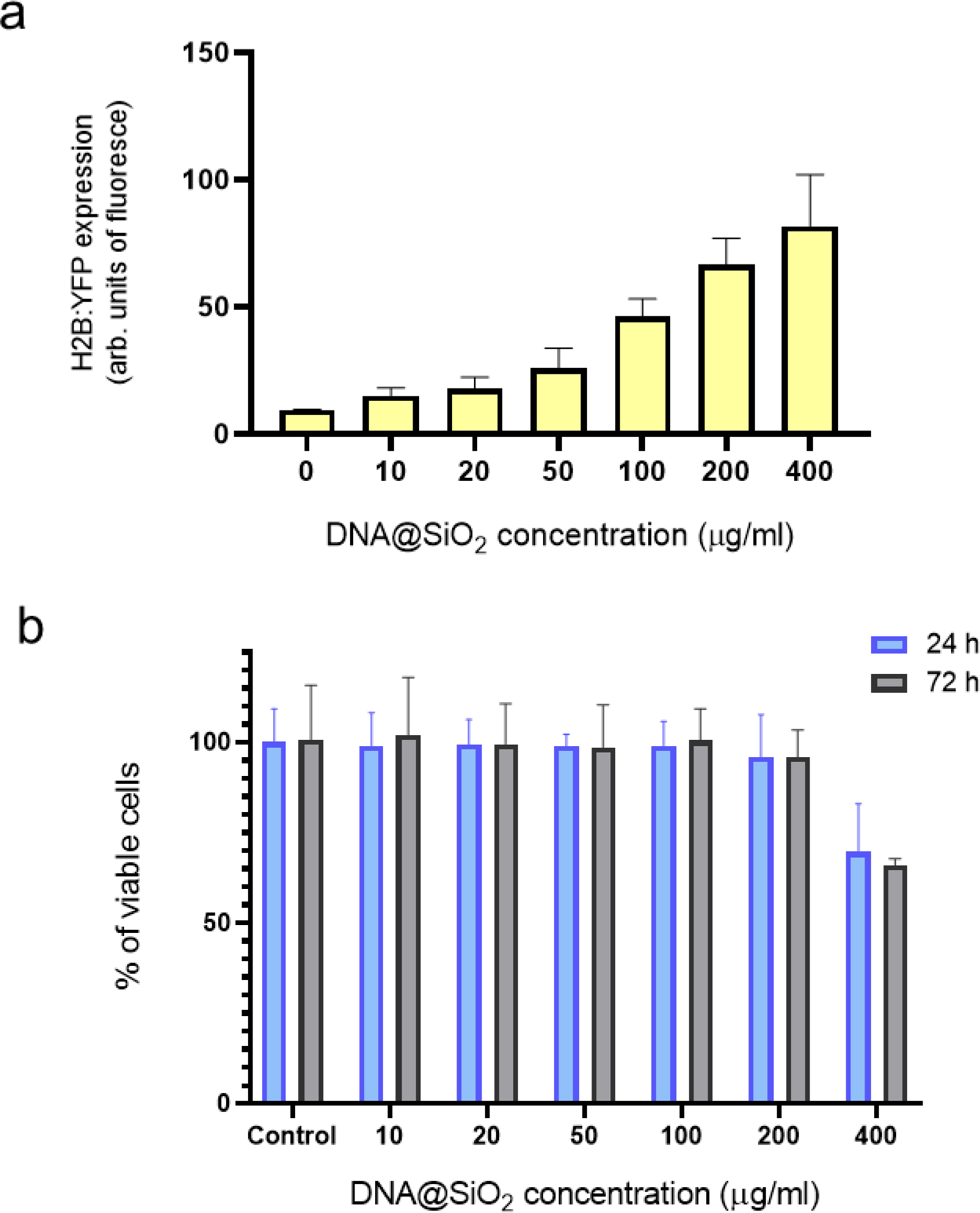
Effect of particle concentration on transfection efficiency and cell viability. (a) Flow cytometry quantification of transfected H2B:YFP protein by using l-DNA#4 particles (mean ± SD, *n* = 3) carrying different DNA@SiO_2_ concentrations (µg/ml IMDM). **(b)** MTT assay showing % of viable cells 24 h and 72 h after l-DNA#4 treatment with the same particles (mean ± SD, *n* = 6).

### 3.5 DNA@SiO_2_ particles are better suited for l-DNA compared to conventional transfection reagents

Next, we wanted to assess how the gene expression efficiency of optimized DNA@SiO_2_ particles loaded with linear DNA (l-DNA) compares to the widely used commercially aviavable Lipofectamine™ 2000. Thus, we transfected HEK 293T cells using both delivery systems. Lipofectamine™ 2000-transfected cells exhibited approximately three times higher gene expression when delivering plasmid DNA compared to the silica gene delivery system (Figure 5). However, the reverse was observed when linear DNA was loaded into the particles, with DNA@SiO_2_ surpassing Lipofectamine™ 2000 transfection efficiency by approximately three-fold. More importantly, linear DNA loaded DNA@SiO_2_ also outperformed Lipofectamine™ 2000 loaded with plasmid DNA by abbut 20% making it the overall supperia transfection system (Figure 5).

**Figure 5.**
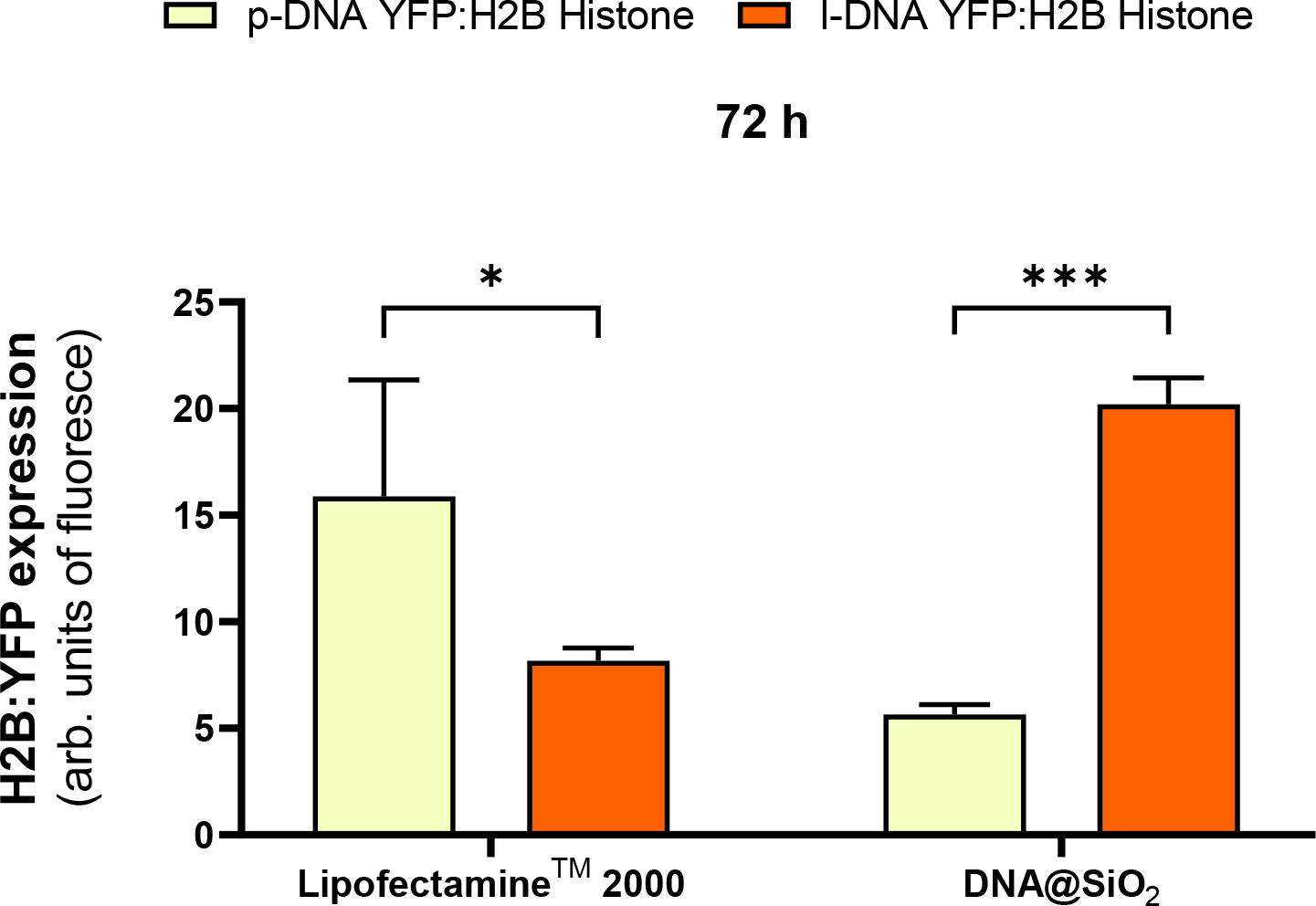
Quantification of H2B:YFP protein expression (mean fluorescence intensity) using Lipofectamine™ 2000 and DNA@SiO_2_ system with p-DNA and l-DNA in HEK 293T cells. Data are shown as the mean ± SD of 3 experimental replicas (*n =* 10,000 cells/replica, *t-*test, \**p* < 0.05, and \*\*\**p* < 0.001).

## 4. Discussion

Transfection of non-circular DNA using conventional non-viral vectors has been limited by their reduced compactability and DNA loading capacity compared to plasmids. Our study presents a novel approach by exploring the use of innovative DNA@SiO_2_ particles, which have already been recognised for their remarkable protection of DNA against temperature, reactive oxygen species (ROS) and nucleases, as well as their ability to sustain gene transfection for up to 14 days. [33] Unlike lipid or polymeric vectors, which typically require compact DNA morphologies for nucleic acid integration, DNA@SiO_2_ particles can encapsulate both plasmidic and linear DNA. Since we have established a maximum loading capacity limit of 4.5 µg DNA per mg of SiO_2_, we used the PCR technique to reduce the DNA molecule size to increase the copies of genes per particle. Using this approach, we could achieve a 16-fold increase in protein expression. This demonstrates the versatility of SiO_2_ nanoparticles to encapsulate DNA regardless of its size and morphology.

When comparing the transfection efficiency of linear DNA (l-DNA) spheres with plasmid DNA (p-DNA) spheres, the protein expression peak occurred at 72 hours after transfection, thus being accelerated compared to the expression profile observed for plasmid DNA which peaked at around 6 days after transfection. We believe that the controlled release of DNA is a result of the outside-in dissolution of compact silica beads, a release mechanism that resulted in faster release of smaller DNA fragments during transfection experiments. Additionally, linear DNA is susceptible to exonucleases and thus might have a shorter lifetime compared to plasmid DNA. In combination with the higher copy number of genes delivered this presumably leads to the observed pattern of higher and earlier peaking gene expression compared to transfection with plasmid DNA. Remakebly, l-DNA#04@SiO_2_ particles are more efficient than standard Lipofectamine™ 2000 in l-DNA as well as p-DNA cell transfection.

While our study used linearised genes (2-4 kb), this delivery method could be adopted to deliver smaller and more diverse nucleic acids, such as oligonucleotides, which are of high interest in therapeutic applications. The results underline the potential application of this system with therapeutic oligonucleotides, as the proposed release mechanism suggests that smaller gene fragments may elicit more robust responses. In addition, the internal entrapment of the genetic cargo opens avenues for additional surface modification of nanoparticles with biomolecules to enhance nanoparticle targeting and biological properties. This would enable targeted gene transfer, avoiding potential problems and side effects associated with viral vectors. Reducing the size of nanoparticles plays a key role, as it allows for improved intercellular dispersion and thus increased cellular uptake. In addition, downsizing opens the door to adding additional layers of silica encapsulating a different gene. This would allow time-controlled release and expression of different genes. [33] For example, this could involve inhibition of disease-related protein expression followed by activation of genes to facilitate protein repair - a promising avenue for therapeutic intervention.

## 5 Conclusion

In this study, we introduce highly efficient gene expression through a novel formulation of amorphous silica nanoparticles as a linear DNA delivery system outperforming LipofectamineTM 2000 in cell transfection efficiency. We demonstrated that small nanoparticle size enhances intercellular and intracellular dispersion, and facilitates controlled release of plasmid and linear DNA templates. These DNA@SiO_2_ particles show promise for gene therapy, offering superior performance in the delivery of linear DNA compared to traditional methods, overcoming size and morphology limitations. Notably, this research would revitalize the role of inorganic vectors, specifically those based in silica, in the storage and delivery of genetic material. Once underestimated, these vectors could be now positioned to exert a considerable influence in gene delivery.

A key highlight of this paper is the newfound accessibility of these nanoparticles, thanks to their straightforward synthesis and composition. This quality makes them applicable across a most of molecular biology laboratories. What sets these particles apart is their capacity to efficiently store linear DNA, such as PCR products, and facilitate their one-step transfection or screening.

## 6 Acknowledgments

MLF acknowledges the financial support from the Spanish Instituto de Salud Carlos iii, and the European Union FEDER funds under Projects ref. PI22/00030, co-funded by the European Regional Development Fund, “Investing in your future” and Grant TED2021-129248 BeI00, the Maria Zambrano Fellowship RMZ-015 funded by MCIN/AEI/10.13039/501100011033 and by the “European Union Next Generation EU/PRTR”. We also thank the Gobierno Regional de Cantabria and IDIVAL for the project Refs IDI 20/22, INNVAL21/19, NEXTVAL 22/12, and IDI-020-022. The figures and graphs have been created with BioRender software (BioRender.com, License ID: 9519A1C8-0002).

