## Supplementary Information for "Biodegradable silica nanoparticles for efficient linear DNA gene delivery"

### Supplementary Figures

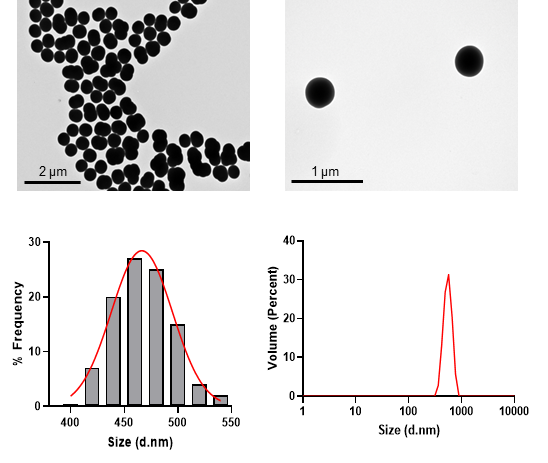

**Figure S1.** TEM characterization and DLS size distribution of p-DNA#1 particles.

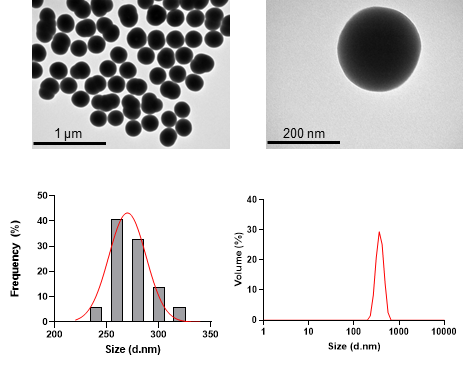

**Figure S2.** TEM characterization and DLS size distribution of p-DNA#2 particles.

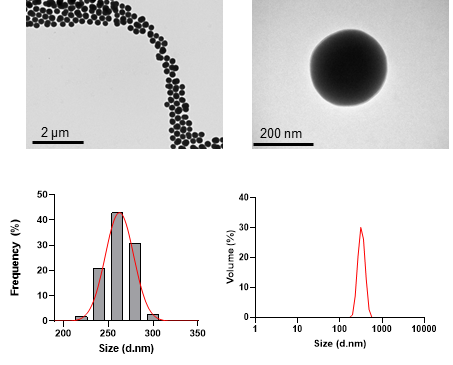

**Figure S3.** TEM characterization and DLS size distribution of p-DNA#3 particles.

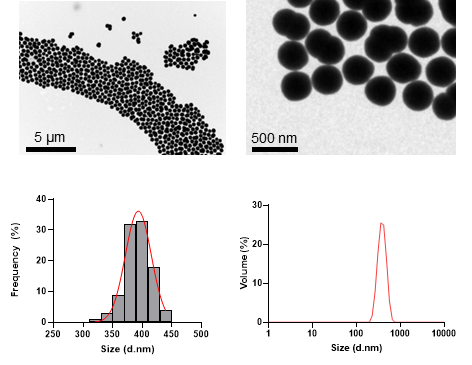

**Figure S4.** TEM characterization and DLS size distribution of p-DNA#4 particles.

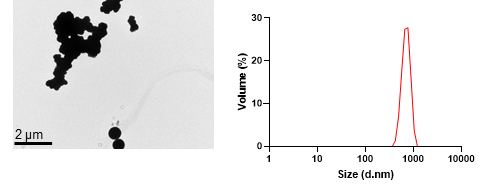

**Figure S5.** TEM characterization and DLS size distribution of p-DNA#5 particles.

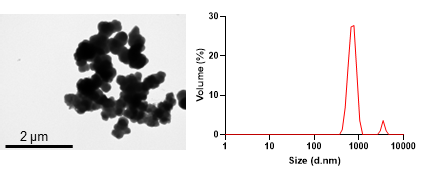

**Figure S6.** TEM characterization and DLS size distribution of p-DNA#6 particles.

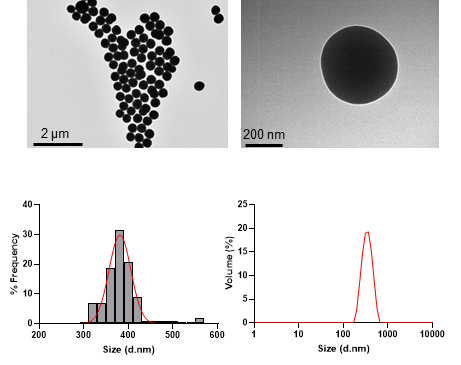

**Figure S7.** TEM characterization and DLS size distribution of l-DNA#1 particles.

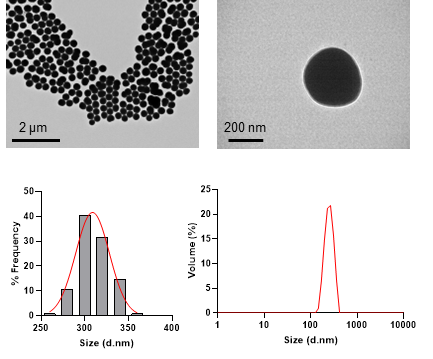

**Figure S8.** TEM characterization and DLS size distribution of l-DNA#2 particles.

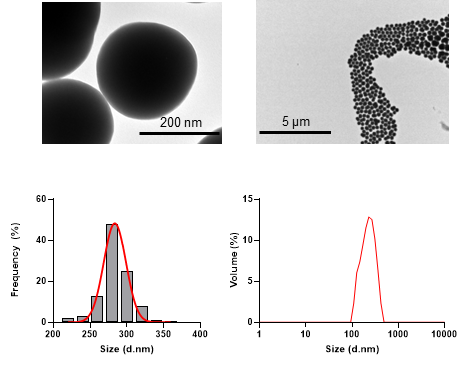

**Figure S9.** TEM characterization and DLS size distribution of l-DNA#3 particles.

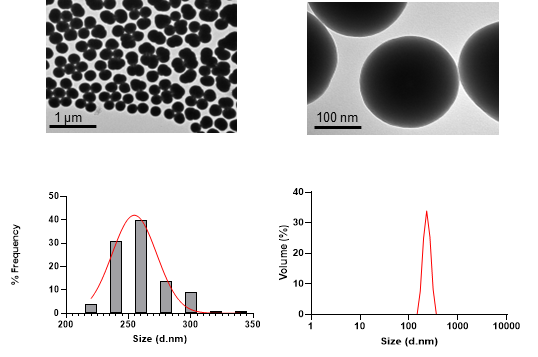

**Figure S10.** TEM characterization and DLS size distribution of l-DNA#4 particles.

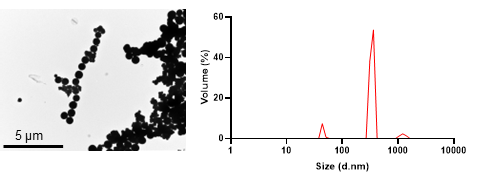

**Figure S11.** TEM characterization and DLS size distribution of l-DNA#5 particles.

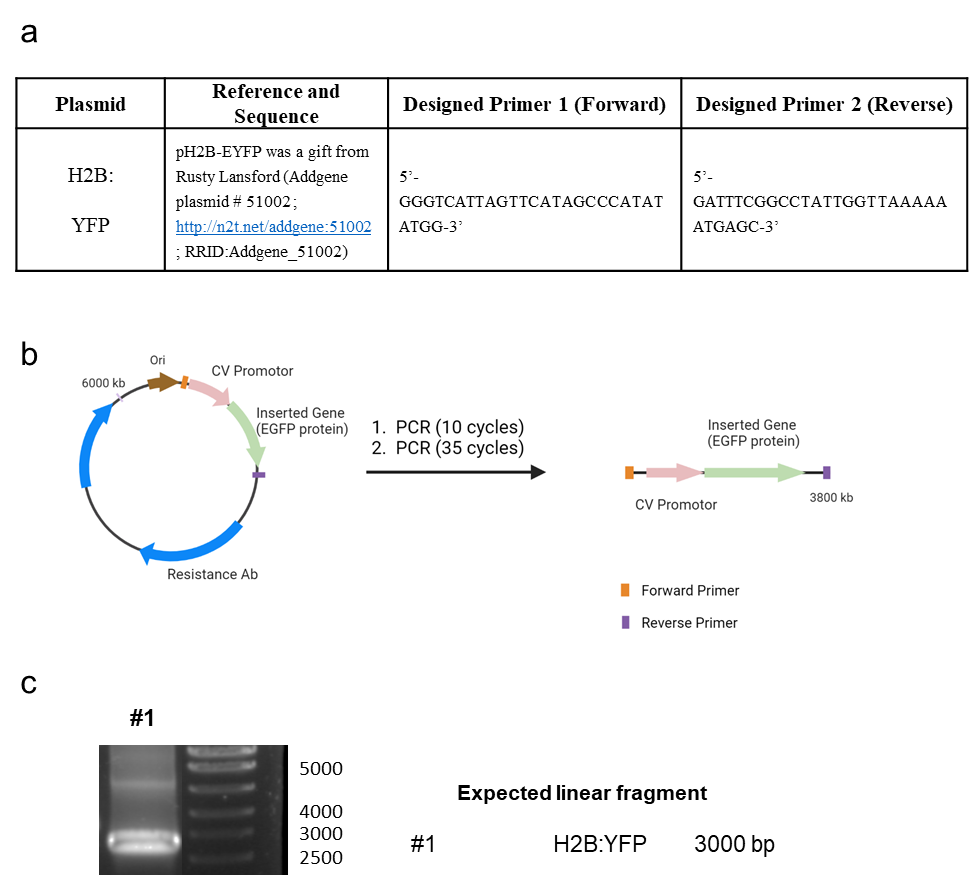

**Figure S12.** p-DNA to l-DNA transformation. **(a)** PCR components and primer sequence design. **(b)** Diagram of the experimental procedure. **(c)** Agarose gels characterization of the synthetized l-DNA.
